# A universal probabilistic spike count model reveals ongoing modulation of neural variability

**DOI:** 10.1101/2021.06.27.450063

**Authors:** David Liu, Máté Lengyel

## Abstract

Neural responses are variable: even under identical experimental conditions, single neuron and population responses typically differ from trial to trial and across time. Recent work has demonstrated that this variability has predictable structure, can be modulated by sensory input and behaviour, and bears critical signatures of the underlying network dynamics and computations. However, current methods for characterising neural variability are primarily geared towards sensory coding in the laboratory: they require trials with repeatable experimental stimuli and behavioural covariates. In addition, they make strong assumptions about the parametric form of variability, rely on assumption-free but data-inefficient histogram-based approaches, or are altogether ill-suited for capturing variability modulation by covariates. Here we present a universal probabilistic spike count model that eliminates these shortcomings. Our method builds on sparse Gaussian processes and can model arbitrary spike count distributions (SCDs) with flexible dependence on observed as well as latent covariates, using scalable variational inference to jointly infer the covariate-to-SCD mappings and latent trajectories in a data efficient way. Without requiring repeatable trials, it can flexibly capture covariate-dependent joint SCDs, and provide interpretable latent causes underlying the statistical dependencies between neurons. We apply the model to recordings from a canonical non-sensory neural population: head direction cells in the mouse. We find that variability in these cells defies a simple parametric relationship with mean spike count as assumed in standard models, its modulation by external covariates can be comparably strong to that of the mean firing rate, and slow low-dimensional latent factors explain away neural correlations. Our approach paves the way to understanding the mechanisms and computations underlying neural variability under naturalistic conditions, beyond the realm of sensory coding with repeatable stimuli.

## 1 Introduction

Classical analyses of neural coding are based on mean spike counts or neural firing rates. Indeed, some of the most paradigmatic examples of the neural code were discovered by regressing neural firing rates to particular sensory stimuli [1, 2] or behavioural covariates [3, 4, 5, 6] to characterize their tuning properties. However, neural spiking is generally not regular. Recordings from many cortical areas show significantly different activity patterns within and across identical trials [7], despite fixing experimentally controlled variables. This irregularity is also seen in continual neural recordings without trial structure [8]. The resulting variability has classically been characterised as ‘Poisson’, with a Fano factor (variance to mean ratio) of one [9], but experimental data also often exhibits significantly more [10, 8, 11, 12] and sometimes less [13, 14] variability, respectively referred to as over- or underdispersion. Moreover, experimental studies have revealed that neural variability generally depends on stimulus input and behaviour [15, 16, 17, 18], and exhibits structured shared variability (‘noise correlations’) across neurons even after conditioning on such covariates. Such correlations can have important consequences for decoding information from neural population activity [19, 20, 21] and reveal key properties of the underlying circuit dynamics [22]. Moreover, theories of neural representations of uncertainty have assigned computational significance to variability as a signature of Bayesian inference [23, 24, 25, 26]. Thus, just as classical tuning curves for firing rates have been crucial for understanding some of the fundamental properties of the neural code, a principled statistical characterisation of neural variability, and its dependence on stimulus and behavioral covariates, is a key step towards understanding the dynamics of neural circuits and the computations they subserve.

The traditional approach to characterising neural variability has been pioneered in sensory areas, and relies on repeatable trial structure with a sufficiently large number of trials using identical stimulus and behavioral correlates [27, 15, 28]. Variability in this case can be quantified by simple summary statistics of spike counts across trials of the same condition. However, this approach does not readily generalise to more naturalistic conditions where covariates cannot be precisely controlled and repeated in an experiment. This more general setting requires statistical methods that take into account temporal variation of covariates for predicting neural count activity. Generalised Linear Models are a popular choice [29], but they only model the dependence of firing rates on covariates – with changes in variability directly coupled to changes in the rate inherent to Poisson spiking. More complex methods for inferring neural tuning [30, 31] and latent structure [32, 33, 34, 35] similarly use restrictive parametric families for spike count distributions, and thus also cannot model changes in variability that are not ‘just’ a consequence of changes in mean counts or firing rates. Conversely, statistical models capable of capturing arbitrary single neuron count statistics, such as histogram-based approaches or copulas [36], do not incorporate dependencies on covariates.

Here we unify these separate approaches, resulting in a single framework for jointly inferring neural tuning, single neuron count statistics, neural correlations, and latent structure. Our semi-parametric approach leads to the universal count model (UCM) for counts ranging from 0 to *K*, in the sense that we can model arbitrary distributions over the joint count space of size (*K* + 1)^*N*^ of *N* neurons. The trade-off between computational overhead and model expressivity is controlled by hyperparameters, with expressivity upper bounded by the true universal model. Our approach extends the idea of a universal binary count model [37] to a finite range of integer counts, while allowing flexible dependence on observed and latent covariates to model non-stationary neural activity and correlations. The flexibility reduces biases from restrictive assumptions in any of the model components. Scalability is maintained by leveraging sparse Gaussian processes [38] with mini-batching [39, 40] to handle the size of modern neural recordings.

We first define the UCM, and then describe how to interpret as well as evaluate model fits. As our model is able to capture arbitrary single neuron statistics, we build on the Kolmogorov-Smirnov test to construct more absolute goodness-of-fit measures. After validating our method on synthetic data that cannot be captured by currently used methods, we apply the model to electrophysiological recordings from two distinct brain regions in mice that show significant tuning to the head direction of the animal [41, 42]. We find that (1) neural activity tends to be less dispersed than common Poisson-like models at higher firing rates, and more dispersed at low rates; (2) mean and variance of counts defy a simple parametric relationship imposed by parametric count distribution families; (3) variability modulation by behaviour can be comparable or even exceed that of the mean count or firing rate; (4) a two-dimensional latent trajectory varying on timescales of ~ 1 s is sufficient to explain away neural correlations but not the non-Poisson nature of single neuron variability. Finally, we discuss related work, limitations and proposed extensions of our model.

## 2 Universal count model

### Notation

Spike count activity of *N* neurons recorded into *T* time bins is formally represented as an N-dimensional time series of non-negative integers. Due to biological constraints, the possible spike counts have some finite upper bound *K*, taken as the highest observed count. We denote probabilities of a spike count distribution (SCD) by a vector ***π*** of length *K* + 1, and use Π to denote the collection of vectors ***π***_*nt*_ for neurons *n* and time steps *t*. Additionally, we denote the count activity by a matrix *Y* ∈ [0, *K*]^*N*×*T*^ with elements *y_nt_*. Input covariates are observed *X* ∈ ℝ^*T*×*D_x_*^ (e.g. animal speed) or latent *Z* ∈ ℝ^*T*×*D_z_*^ (to capture e.g. attention), with range depending on topology [43]. We denote their elements *x_td_* and *z_tq_* with observed and latent dimension *d* and *q*, respectively.

### 2.1 Generative model

The big picture is to model counts *Y* with dependence on *X*. For each neuron, our model consists of *C* Gaussian process (GP) priors, a basis expansion 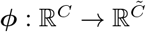, and a linear-softmax mapping

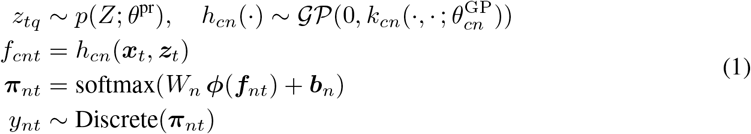

where 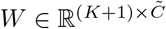, ***b*** ∈ ℝ^*K*+1^, and the GP covariance functions *k_cn_* have hyperparameters 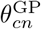. The use of non-parametric GP mappings with point estimates for *W* and ***b*** leads to a semi-parametric model with parameters *θ*, see details in Appendix E. The overall generative model *P_θ_*(*Y*|*X*) is depicted schematically in Figure 1. Note the model specifies a prior *p*(Π|*X*) over joint SCDs, conceptually similar to Dirichlet priors [37] but allowing non-parametric dependence on *X*. With latent input *Z*, our model can flexibly describe multivariate dependencies in joint SCDs as conditional independence across neurons no longer holds when marginalizing over *Z* [44]. In addition, *p*(*Z*) models temporal correlations in the latent states. We use Markovian priors (details in subsection E.2)

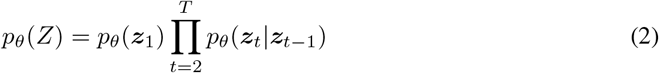

with *θ*^pr^ absorbed into model parameters *θ* for compactness. This allows the model to flexibly capture both neural and temporal correlations in Y. To attain scalability, we use sparse GPs [38].

**Figure 1:**
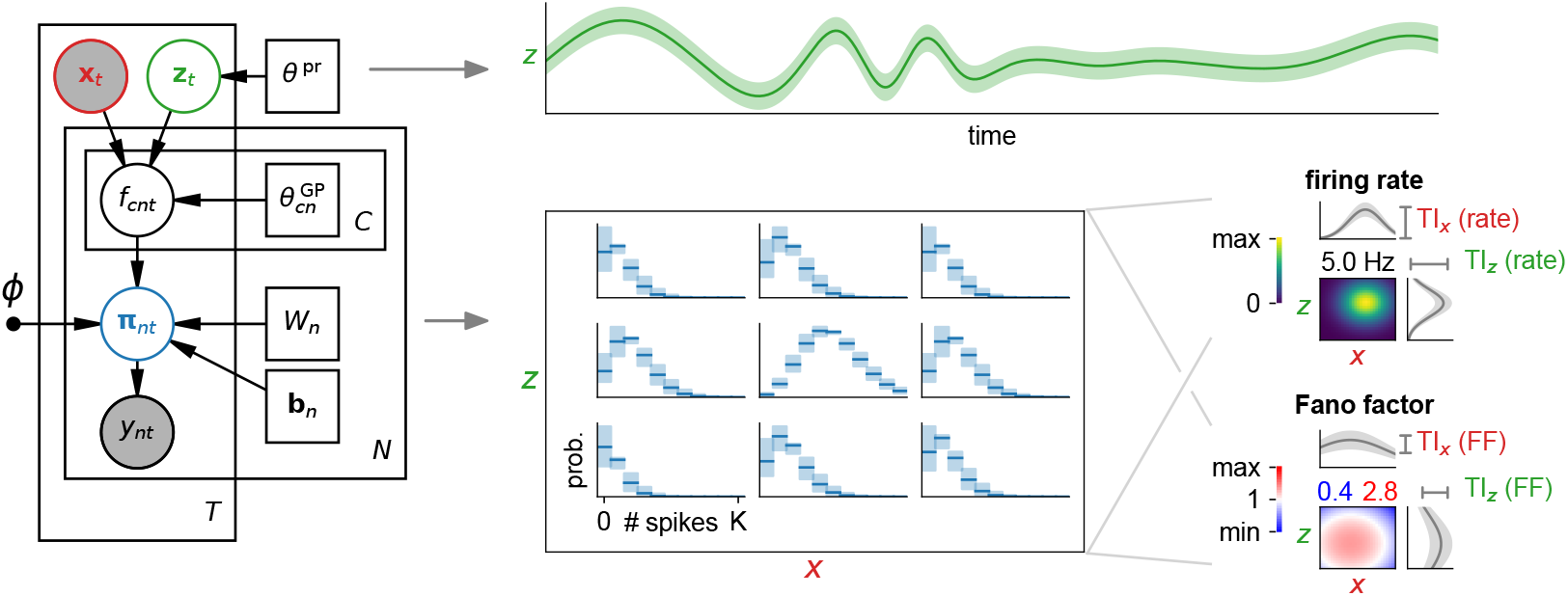
Schematic of the UCM and the workflow. Left: graphical model corresponding to Equation 1, with shaded circles as observed, open circles as latent, and squares as deterministic variables. Filled dots represent fixed quantities. Middle: example inference of model posterior Equation 3, with inferred latent trajectories (green, top) and covariate-dependent SCDs (blue, bottom) that depend on both observed *x* and latent *z* covariates. Note we only show the posterior over a single SCD evaluated on a (*x,z*) grid, whereas the full posterior defines SCDs over all neurons. Right: obtaining interpretable spike count statistics from the SCDs (see subsection 2.3). Examples show firing rate and Fano factor tuning curves over observed *x* and latent *z* covariates, either jointly (heatmaps) or marginalized (grey curves). The depth of modulation in marginalized tuning curves is used to extract a tuning index (TI) for the chosen subsets of covariates, see Equation 6.

Depending on *C* and basis functions ***ϕ***(·), we obtain an approximation to the true universal prior on joint SCDs, with ‘universal’ referring to the ability to capture any joint SCD over all neurons. Arbitrary single neuron statistics can be captured when *C* = *K* with ***ϕ***(***f***) = ***f***, but fitting is computationally expensive when *N* × *C* ≫ 1. For capturing all correlations, the model also requires a sufficiently large latent space. One controls the trade-off between model expressiveness and computational overhead through *C* and ***ϕ***. Larger expansions ***ϕ*** allow one to model count distributions more expressively with small *C*, e.g. the element-wise linear-exponential ***ϕ***(***f***) = (*f*_1_, *e*^*f*_1_^,…, *f_C_*, *e*^*f_C_*^) covers a range of distributions including the truncated Poisson with only *C* = 1 (see subsection A.3).

### 2.2 Stochastic variational inference and learning

For the joint model distribution *p_θ_* (*Y*, Π, *Z*|*X*) = *P*(*Y* |Π) *p_θ_*(Π|*X, Z*) *p_θ_* (*Z*), with count distributions *P*(*Y*|Π), we approximate the posterior by *q_θ,χ,φ_*(Π, *Z*|*X*) that factorizes in the form

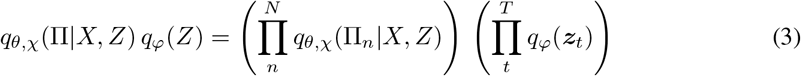

with *φ* and *χ* the variational parameters for latent states and the sparse Gaussian process posterior (Appendix E), respectively. Note that we use a factorized normal *q*(*Z*) for Euclidean *Z*, and a wrapped normal for circular *Z* based on the framework of reparameterized Lie groups [45, 43]. The posterior over count probabilities *q_θ,χ_*(Π|*X, Z*) is defined as mapping the sparse Gaussian process posterior *q_θχ_*(*F*|*X, Z*) through Π(*F*) (Equation 1), a deterministic mapping. This is analytically intractable, so in practice it is represented by Monte Carlo samples. Using stochastic variational inference [46], we minimize an upper bound on the negative log marginal likelihood

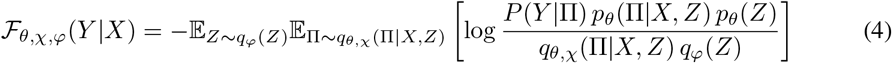

known as the variational free energy. This objective leads to tractable terms (subsection E.1), allowing us to infer the approximate posterior as well as a lower bound of the log marginal likelihood [47, 40]. We use Adam [48] for optimization, see details of implementation and model fitting in Appendix E.

### 2.3 Obtaining interpretable spike count statistics from the model

#### Characterizing spike count distributions

From the posterior *q*(Π|*X*)^1^, we can compute samples of the posterior of any statistic of spike counts as a function of covariates. Single neuron statistics in particular can be characterized by tuning curves for both mean firing rates and Fano factors (FF)

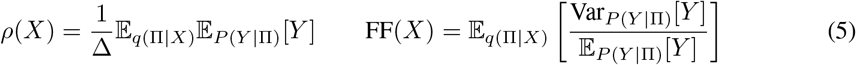

with time bin length Δ. The model also quantifies private neuron variability that cannot be explained away by regressing to shared input (both observed and latent) through *P_n_*(*y_tn_*|***x***_*t*_).

To quantify the sensitivity of a some aspect of neuron activity to a set of covariates ***x***\*, we define a tuning index (TI) with respect to a count statistic *T_y_*(***x***\*)

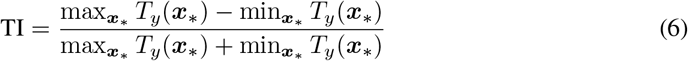

with *T_y_* (***x***\*) evaluated under the mean posterior SCD marginalized over all other covariate dimensions complementary to ***x***\*. These marginalized distributions are computed using observed input ***x***_*t*_ (subsection D.5). Resulting marginalized tuning curves for TIs are depicted conceptually in Figure 1.

#### Generalized *Z*-scores and noise correlations

The deviation of activity from the predicted statistics is commonly quantified through *Z*-scores [8, 49, 17], which are computed as 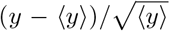 with 〈*y*〉 being the mean count in some time bin. If neural activity follows a Poisson distribution, the distribution of *Z*-scores asymptotically tends to a unit normal when average counts 〈*y*〈 ≫ 1 (Appendix B). To generalize the normality of the *Z*-score for arbitrary counts and SCDs, we use

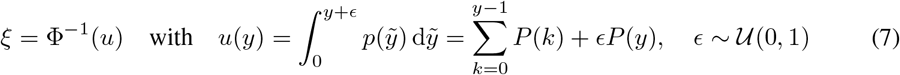

with Φ(·) the unit Gaussian cdf., and dequantization noise *ϵ* to get continuous quantiles 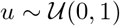 and generalized *Z*-scores 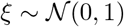 from the probability integral transform.

With *ξ*, one can completely describe single neuron statistics with respect to the model. Correlations in the neural activity however will cause to be correlated. We define generalized lagged correlations *r_ij_* (Δ) ∈ [−1,1] and Fisher *Z* ∈ ℝ that is more convenient for statistical testing

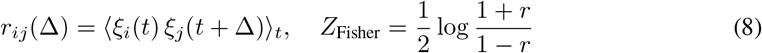

which describes spatio-temporal correlations not captured by the model. Noise correlations [50] refer to the case of Δ = 0, when *r_ij_* becomes symmetric.

### 2.4 Assessing model fit

Our model depends on a hyperparameter *C* ≤ *K* that trades off flexibility with computational burden. In practice, one likely captures the neural activity accurately with *C* well below *K* and a simple basis expansion as the linear-exponential above or quadratic-exponential 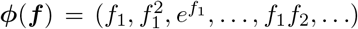. This can be quantified by the statistical measures provided below, and allows us to select appropriate hyperparameters to capture the data sufficiently well.

To assess the model fit to neural spike count data, a conventional machine learning approach is to evaluate the expected log-likelihood of the posterior predictive distribution on held-out data *Y*, leading to the cross-validated log-likelihood

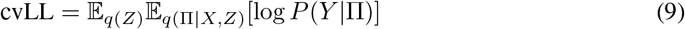

where we cross-validate over the neuron dimension by using the majority of neurons to infer the latent states *q*(*Z*) in the held-out segment of the data, and then evaluate Equation 9 over the remaining neurons. Without latent variables, we simply take the expectation with respect to *q*(Π|*X*). However, the cvLL does not reveal how well the data is described by the model in an absolute sense. Likelihood bootstrap methods are possible [28], but become cumbersome for large datasets. To assess whether the neural data is statistically distinguishable from the single neuron statistics predicted by the model, we use the Kolmogorov-Smirnov framework [51] with *u* from Equation 7 across time steps *t*

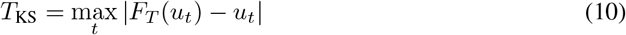

with empirical distribution function *F_T_*(*u*) (for details see Appendix B). This scalar number is positive and does not indicate whether the data is under- or overdispersed relative to the model. For this, a useful measure of dispersion is the logarithm of the variance of *ξ* with a correction

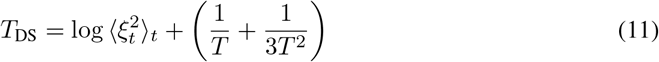

which provides a real number with positive and negative sign indicating over- and underdispersion, respectively. Its sampling distribution under 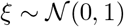 is asymptotically normal, centered around 0 (due to the additive parenthetical term) with a variance depending on the number of timesteps *T* (subsection B.3). This extends the notion of over- and underdispersion beyond the usual definition of dispersion relative to Poisson models [52]. To quantify whether the model has captured noise correlations in the data, we compute with respect to the mean posterior predictive distribution

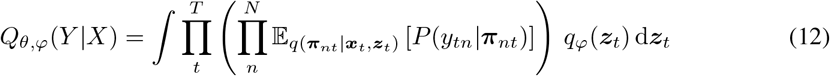

which whitens *ξ*, hence reducing noise correlations *r* in Equation 8, if correlations are explained away by co-modulation of neurons due to shared low-dimensional factors [44]. These are captured through latent states *Z* using the posterior *q_φ_*(*Z*), inferred from the same data used to compute *ξ*. Overall, this can be interpreted as treating *Z* as if it was part of the observed input to the model.

## 3 Results

In the following, we use *C* = 3 with an element-wise linear-exponential basis expansion (subsection 2.1). This empirically provided sufficient model flexibility as seen in goodness-of-fit metrics. We use an RBF kernel with cosine distances in case of angular input dimensions (subsection E.3).

### 3.1 Synthetic data

Animals maintain an internal estimate of their head direction [4, 42, 53]. Here, we extend simple statistical models of head direction cell populations [54] for validating the ability of the UCM to capture complex count statistics, as well as neural correlations through latent structure. The task is to jointly recover the ground truth count likelihoods, their tuning to covariates, and latent trajectories if relevant from activity generated using two synthetic populations. The first population was generated with a parametric heteroscedastic Conway-Maxwell-Poisson (CMP) model [55], which has decoupled mean and variance modulation as well as simultaneously over- and underdispersed activity (Fano factors above and below 1). The second population consists of Poisson neurons tuned to head direction and an additional hidden signal, which gives rise to apparent overdispersion [28] as well as noise correlations when only regressing to observed covariates. For mathematical details of the count distributions and synthetic populations, see Appendix A and Appendix D, respectively.

We compare our UCM to the Poisson GP model [33] and the heteroscedastic negative binomial GP model, a non-parametric extension of [55]. To show data-efficiency and regularization benefits of GPs, we also compare to a UCM with an artificial neural network (ANN) mapping replacing the GP mapping. For details of the baseline models, see Appendix D. For cross-validation we split the data into 10 roughly equal non-overlapping segments, and validated on 3 chosen segments that were evenly spread out across the data. When a latent space was present, we used 90% of the neurons to infer the latent signal while validating on the remaining neurons, and repeated this for non-overlapping subsets. We rescale the log-likelihoods by the ratio of total neurons to neurons in subset and then take the average over all subsets to obtain comparable cross-validation runs to regression.

Figure 2A shows that only the UCM successfully captures the heteroscedastic CMP data, indicated by *T*_KS_. Baseline models cannot capture frequent cases where the Fano factor drops below 1. In addition, we observe that using a Bayesian GP over an ANN mapping in the model leads to a reduction in overfitting, especially in the latent setting (Figure 2A). Therefore, all other analyses with the UCM reported here used the GP mapping. Figure 2B shows that the modulated Poisson population activity is seen by a Poisson regression model as overdispersed, indicated by *T*_DS_. Our model flexibly captures the overdispersed single neuron statistics, independent from noise correlations *r_ij_* that are captured when we introduce a Euclidean latent dimension. As expected, the *ξ* scatter plots shows whitening under the posterior predictive distribution when the correlations are captured.

**Figure 2:**
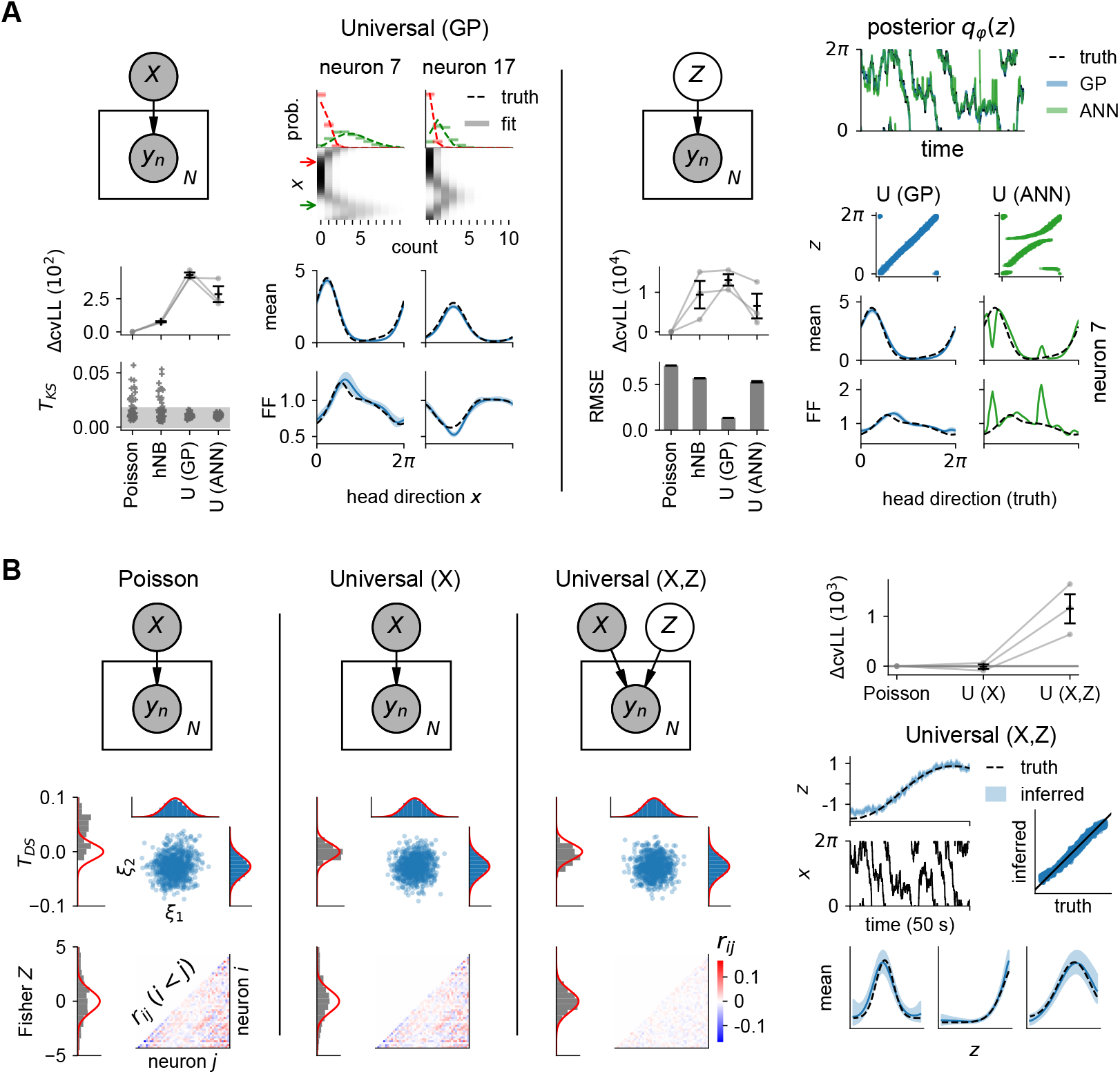
Model validation with two synthetic head direction cell populations. **(A)** Applying Poisson, heteroscedastic negative binomial (hNB) and universal (U, with either GP or ANN mappings) models to synthetic data from the heteroscedastic CMP population. Error bars indicate s.e.m. over cross-validation runs. Shaded regions for tuning curves and *T*_KS_ indicate the 95% CI. ΔcvLL is the difference w.r.t. Poisson baseline. Left: regression models, visualizing SCDs (top right), tuning curves (bottom right), and fitting scores (bottom left) for two representative cells. Right: models with an angular latent variable, visualizing inferred latent states (top right), comparing the GP to ANN model (bottom right), and plotting fitting scores (bottom left). Root-mean-squared errors (RMSE) between the inferred latent and ground truth uses the geodesic distance on the ring (subsection D.9). **(B)** Applying regression (Poisson, Universal (X)) and joint latent-observed (Universal (X,Z)) models to the modulated Poisson population data. Left: three columns showing progressively how single neurons variability, with *ξ* and *T*_DS_ (middle), and noise correlations, with *r_ij_* and Fisher *Z* (bottom), are captured (see subsection 2.4). Right: ΔcvLL for all models (top) and visualization of the joint latent-observed model input spaces (middle) and tuning curves (bottom).

### 3.2 Mouse head direction cells

We apply the UCM to a recording of 33 head direction cells in the anterodorsal thalamic nucleus (ADn) and the postsubiculum (PoS) of freely moving mice [41, 42], see subsection D.2. Neural data was binned into 40 ms intervals, giving *K* = 11. Note that observed count statistics differ with bin size, see subsection C.2 for a discussion. Regression was performed against head direction (HD), angular head velocity (AHV), animal speed, two-dimensional position in arena, and absolute time, which collectively form *X* in this model. We used 64 inducing points for regression, and added 8 for every latent dimension added (Appendix E). Cross-validation was performed as in synthetic experiments, but with 6 validation segments and subsets of 85% of neurons to infer the latent signal.

Figure 3A shows that for regression hNB overfits and performs worst, despite containing Poisson as a special case. However, this limit is not reached in practice due to the numerical implementation, see Appendix A. Only the UCM captures the training data satisfactorily with respect to confidence bounds for *T*_KS_ and *T*_DS_, although the data remained slightly underdispersed to the model with *T*_DS_ values slightly skewed to negative. Compared to the Poisson model, the cvLL is only slightly higher for the UCM as the data deviates from Poisson statistics in subtle ways. We see both FF above and below 1 (over- and underdispersed) across the neural firing range in Figure 3B, with quite some neurons crossing 1. Correspondingly, FF-mean correlations coefficients are often negative. Their spread away from ±1 indicates firing rate and FF do not generally satisfy a simple relationship, especially for examples such as cell 27. Furthermore, ADn neurons seem to deviate less from Poisson statistics. From Figure 3C, we note in particular that FFs tend to decrease at the preferred head direction, but rise transiently as the head direction approaches the preferred value. We also see that tuning to speed and time primarily modulates variability rather than firing rates. All of this is impossible to pick up with baseline models, which constrain FF ≥ 1 as well as FF increasing with firing rate (Appendix A). Finally, we see more tuning of the firing rate to position in PoS cells.

**Figure 3:**
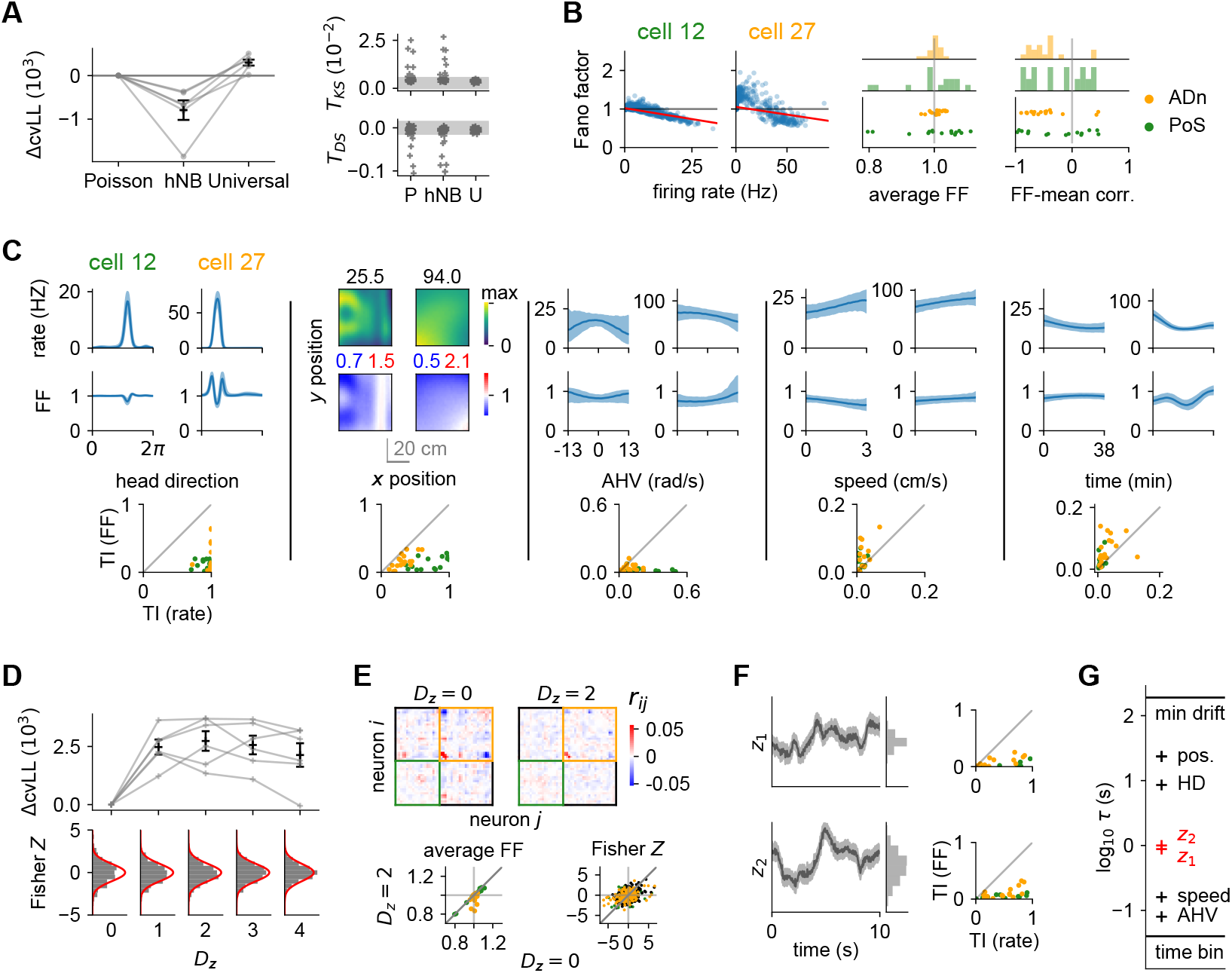
Application to mouse head direction cells in the anterodorsal thalamic nucleus (ADn) and the postsubiculum (PoS). **(A)** Goodness-of-fit for Poisson (P), heteroscedastic negative binomial (hNB) and universal (U) regression with covariates described in the text. Our method (U) outperforms baselines (*p* = 7 · 10^-3^ for ΔcvLL w.r.t. Poisson, one-sample *t*-test). **(B)** Left: Fano factor (FF) versus mean count of the predictive distribution for all time steps (scatter dots) for two representative cells. Red lines are linear regression fits to scatter points. Right: average FF across time steps and Pearson *r* correlation between FF and mean count for all cells, with colour indicating region ADn (orange) or PoS (green). **(C)** Top: conditional tuning curves (subsection D.5) of the UCM with complementary ***x*** at preferred HD, positioned at centre of arena, with zero speed, AHV and at time *t* = 0. Bottom: TIs w.r.t. corresponding covariates for the FF and firing rates. **(D)** Adding Euclidean latent dimensions *D_z_*. Top: ΔcvLL w.r.t. regression only. Bottom: Fisher *Z* histogram, with red curves its sampling distribution. **(E)** Comparison between models with latent space dimensions *D_z_* = 2 and *D_z_* = 0. Top: off-diagonal noise correlations *r_ij_*, with neurons ordered by region first (PoS and ADn), and then by preferred head direction within region. Bottom: average FF (same as in (B)) and Fisher *Z* under the posterior predictive distribution Equation 12. **(F)** Left: inferred latent trajectories for *D_z_* = 2. Right: corresponding TIs for FF and firing rates. **(G)** Time scales for covariates computed from autocorrelograms (Appendix D). Note pos. refers to both *x* and *y* position (near-identical values). The top horizontal line is the minimum kernel length scale over absolute time across neurons. The bottom line depicts the 40 ms time bin. Error bars in **A** and **D** show s.e.m. over cross-validation runs. Shaded regions for *T*_KS_, *T*_DS_, tuning curves and latent states show the 95% CI.

When adding latent dimensions, Figure 3D shows a peak in the cvLL at two dimensions, where correspondingly the Fisher Z distribution starts to match the unit normal well. Kernel length scales however did not indicate redundant latent subspaces for higher dimensions as expected for automatic relevance determination, likely due to mixing of latent dimensions. Notice the noise correlation patterns in Figure 3E tend to show positive correlations for similarly tuned neurons roughly around the diagonal of blocks, as expected from ring attractor models [22]. Intrinsic neuron variability, roughly quantified by the average FF, further decreased and thus become even more underdispersed when considering additional tuning to latents, in particular for ADn. In addition, latent signals primarily modulate firing rate as seen from TIs in Figure 3F. When looking at time scales of covariates in Figure 3G (computed as the decay time constant of the autocorrelogram (Appendix D), the latent processes seem to vary on time scales right in the gap of behavioural time scales.

Tangential to our main contribution of characterising the detailed structure of neural variability, our results have another novel element that does not specifically rely on the UCM. Using GP-based non-parametric methods, we successfully estimated the tuning of cells to as many as 8 different covariates (6 observed + 2 latent, see Figure 3G) in a statistically sound fashion, while previous GP-based approaches typically only consider around 2 to 3 input dimensions [56, 33, 43]. Specifically, one of our covariates was absolute experimental time to capture non-stationarities in neural tuning. As a result, our model captured several experimental phenomena that are studied separately in the literature: drifting neural representations [57, 58, 59], anticipatory time intervals [54] and conjunctive tuning to behaviour [60]. We also applied the model in a purely latent setting similar to the example in Figure 2A, with the UCM uncovering a latent signal more closely correlated to the head direction compared to baseline models. These additional results are presented in Appendix C.

## 4 Discussion

### Related work

Neural encoding model provide a statistical description of neural count activity, and typically rely on a parametric count likelihood, such as Poisson [33] or negative binomial [30, 31, 61], that is often mismatched to empirical count statistics. Heteroscedastic versions additionally regress the dispersion parameter of count distributions to covariates [62, 63]. This has shown improvements in stimulus decoding and more calibrated posterior uncertainties [55]. Copula-based models [36, 64] separate marginal distributions of single neurons from the multivariate dependency structure parameterized by the copula family, and thus remove parametric constraints on single neuron count statistics. Generally, the idea of a universal model that can capture arbitrary joint distributions has been explored for binary spike trains [37]. However, neither approach naturally incorporates modulation of spike count distributions by input covariates.

Our model deals with discrete spike counts ranging from 0 to K in a manner similar to categorical output variables in classification, where existing GP-based approaches pass function points directly through a softmax nonlinearity [65, 66]. Our approach instead relies on a basis expansion and linear-softmax mapping. At small time bins where *K* = 1, our model becomes identical to Bernoulli models [67] and comparable to point process models [51, 68, 69, 70]. In these cases, modulation of count variability becomes inseparable from firing rate modulation, making it difficult to generalize for heteroscedasticity in an interpretable manner and thus inconveniet for studying response variability. Introducing unobserved input variables incorporates aspects of Gaussian process latent variable models [71, 72]. Such models have been applied to neural data to perform dimensionality reduction [33], with extensions to non-Euclidean latent spaces and non-reversible temporal priors [43, 73].

### Limitations and further work

The empirical choice of hyperparameters *C* and basis functions *ϕ* is based on achieving sufficient model flexibility, as confirmed with the Kolmogorov-Smirnov approach. Recently, a multivariate extension of the Kolmogorov-Smirnov test has been proposed to directly test multivariate samples against the model [74], instead of looking at single neuron statistics. Alternatively, one could perform ARD [75, 76, 61] by placing a Gaussian prior on *W*, allowing automatic selection of relevant dimensions once a basis expansion is chosen. Another avenue for future work could consider going completely non-parametric by adding a count dimension to the input space, which is evaluated at counts 0 to K for every time step. This however increases the number of evaluation points by a factor *K* +1. For high-dimensional input stimuli common in sensory areas, deep kernels [77] provide a scalable modification of our framework. In addition, extending our model with more powerful priors for latent covariates, such as Gaussian process priors [33, 73], can improve latent variable analysis, especially at smaller time bins where the temporal prior influence becomes more important. Regularization methods may help to decorrelate inferred trajectories [78, 79].

### Conclusion and impact

We introduced a universal probabilistic encoding model for neural spike count data. Our model flexibly captures both single neuron count statistics and their modulation by covariates. By adding latent variables, one can additionally capture neural correlations with potentially interpretable unobserved signals underlying the neural activity. We applied our model to mouse head direction cells and found count statistics that cannot be captured with current methods. Neural activity tends to be less variable at higher firing rates, with many cells showing both over- and underdispersion. Fano factors and mean counts generally do not show a simple relation and can even be decoupled, with Fano factor modulation comparable or in some cases even exceeding that of the rate. Finally, we found that a two-dimensional latent trajectory with a timescale of around a second explained away noise correlations in these cells.

Neural variability is usually not considered on the same footing as mean firing rates, with models assigning most computational relevance to rates [80, 81]. However, recent work on V1 has started to explore variability as playing a computationally well-defined useful role in the representation of uncertainty [24, 25, 22, 26]. The framework introduced in this paper provides a principled tool for empirically characterising neural variability and its modulations – without the biases inherent in traditional approaches, which would likely miss potentially meaningful patterns in neural activities beyond mean rates. Our model has the potential to reveal new aspects of neural coding, and may find practical applications in designing and improving algorithms for brain-machine interfaces. As progress is made in scaling and applying such technology beyond research environments [82], it becomes increasingly more important to maintain transparency, e.g. through open source code, and to raise awareness of potential ethical issues [83].

## Acknowledgments and Disclosure of Funding

This work was supported by the Cambridge European and Wolfson College Scholarship by the Cambridge Trust (D.L.) and by the Wellcome Trust (Investigator Award in Science 212262/Z/18/Z to M.L). We are grateful to K.T. Jensen and A. Melkonyan for helpful feedback on the manuscript.

## Supplementary Material

### A Parametric count distributions

#### A.1 Poisson distribution

The Poisson count distribution is defined with a mean count λ

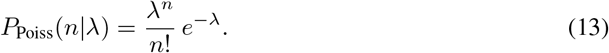

where 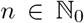. It describes a process where discrete events arriving in a time window are all independent of each other. Mathematically, this is consistent with Equation 13 being the limit of a binomial distribution *P*_Bin_(*n, p*) with *n* → ∞ and *p* → 0, such that *N_p_* = λ.

The Poisson distribution is characterized by the equality of its mean and variance, leading to a Fano factor 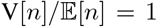. Another convenient property is that the sum of two independent Poisson processes is itself a Poisson process with λ = λ_1_ + λ_2_. This can be shown directly by considering 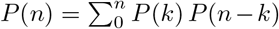 or casting it as a limit of the sum of two Bernoulli processes, and follows intuitively from the fact that spike times are independent of each other. As a consequence, for the inhomogeneous case where we have a time-dependent rate λ(*t*) the count distribution over a longer interval is still Poisson with average 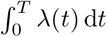. Note that this property does generally not hold for non-Poisson distributions, where the count distribution of a sum of counts in separate time windows is not related to the original count distribution in a simple way.

#### A.2 Non-Poisson count distributions

To account for over- and underdispersed neural activity in real data, i.e. Fano factors above and below 1, other distributions than the Poisson count distribution have been used, and we present common families below.

##### A.2.1 Zero-inflated Poisson

A common way to introduce overdispersion is to model excess zero counts, which in this context leads to the zero-inflated Poisson (ZIP) process [1]. The count distribution is given by

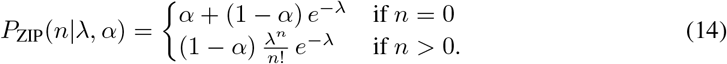

The parameterization leads to 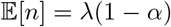 and *V*[*n*] = λ(1 – *α*) + λ^2^*α*(1 – *α*) using the law of total variance.

##### A.2.2 Modulated Poisson distributions

One perspective of non-Poisson distributions is that they arise from noise in the rate parameters λ. Such count processes are referred to as modulated Poisson processes. From a probabilistic point of view, the resulting count distribution is a marginalization

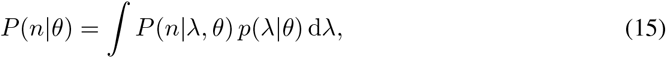

with noise parameters *θ*. A recently proposed flexible spike count model that can give rise to different mean-variance relationships, including decreasing Fano factors at high firing rates similar to what is observed in Figure 4B, builds on this framework [2]. However, the modulated Poisson process can only account for overdispersion with respect to the base Poisson process. Adding noise cannot lead less variability here, and this implies that Fano factors are bounded from below by 1.

**Figure 4:**
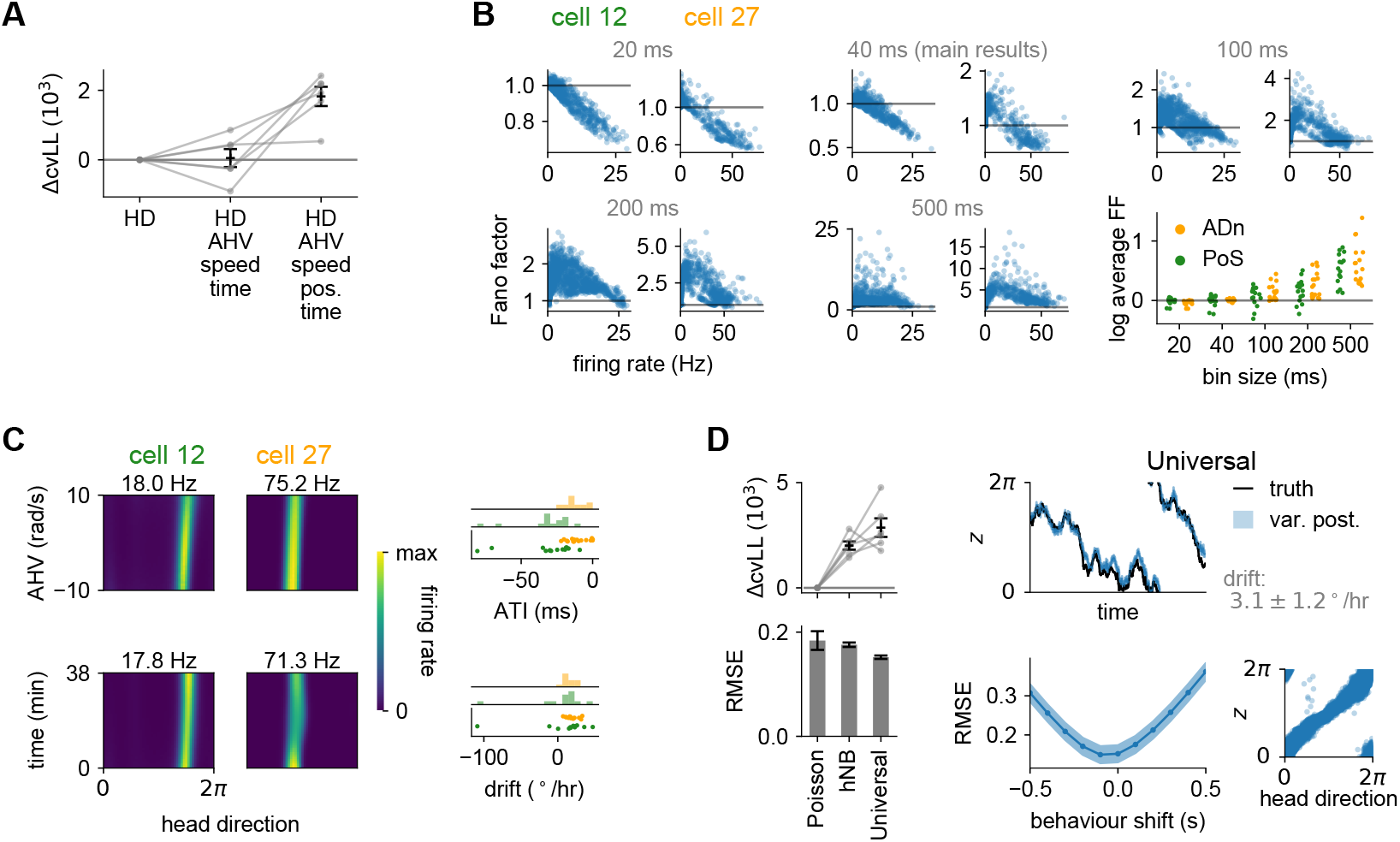
Additional analysis on head direction data. **(A)** Model comparison of the UCM with different regressors (HD: head direction, AHV: angular head velocity, speed: running speed, pos.: animal position, time: experimental time). ΔcvLL shows cross-validated log-likelihood w.r.t. the model with only HD as the regressor. **(B)** Fano factor and mean counts as predicted by the mean posterior count distribution with each dot representing one time step, similar to Figure 3B. We show examples for two representative cells, from model fits at various bin sizes (20, 40, 100, 200 and 500 ms). On the bottom right, we plot the log of the mean Fano factor across all time steps against the bin size for all cells in the data recorded in ADn (orange) and PoS (green). **(C)** Left: Joint conditional tuning curves of firing rate in the full regression model (with complementary covariates fixed at the same values as in Figure 3C) as a function of AHV-HD (top) and time-HD (bottom) show two distinct experimental phenomena: anticipatory tuning and neural representational drift, respectively, for two representative cells. Right: anticipatory time intervals (ATI, top) and drifts (bottom) as defined in subsection C.3, with corresponding histograms, for selected cells (subsection D.8) in the data from ADn (orange) and PoS (green). **(D)** Application of the UCM with only latent regressors. Left: ΔcvLL with respect to Poisson baseline (top) and root-mean-squared error (RMSE, bottom) of estimated latent variable w.r.t. true head direction for different variants, using a Poisson, heteroscedastic negative binomial (hNB), and universal likelihood. Our model (Universal) performs best (*p* =1.3 · 10^-3^ for ΔcvLL, one-sample *t*-test). Right: experimentally observed head direction (black) and estimated latent variable (blue; shaded region shows the 95% CI) in the fitted UCM model as a function of time (top left) and compared against each other (bottom right), matched by fitting a constant angular offset, a sign reversal, and linear drift in time (top right, grey; see Equation 45). We also temporally shifted observed head direction (behaviour) w.r.t. the latent signal, and repeated the same fitting procedure to compute the cross-validated RMSE (bottom left, shaded region shows s.e.m. over cross-validation runs). Error bars in **A** and **D** show s.e.m. over cross-validation runs.

##### A.2.3 Negative binomial

The negative binomial distribution is based on independent Bernoulli trials like the binomial distribution. However, now we count the number of successes before *r* failures are observed. If we have Bernoulli trials with success probability *p*, one can obtain the negative binomial distribution with parameterization 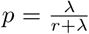

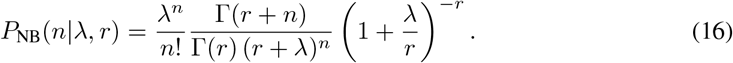

Note this distribution is a specific instance of a modulated Poisson process (Equation 15), with 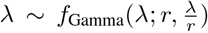. The parameterization is such that 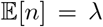 holds, but 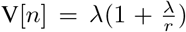 making it overdispersed with respect to a Poisson distribution. In practice, numerical evaluation of the Poisson limit when *r* = 0 is only approximate due to the numerical precision of the relevant function implementations.

##### A.2.4 Conway-Maxwell-Poisson

A distribution that handles both over- and underdispersed count distributions is the Conway-Maxwell-Poisson distribution [3]

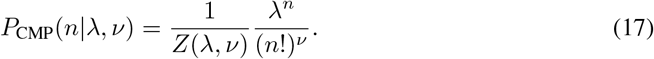

The normalization constant has no closed form expression and must be evaluated numerically

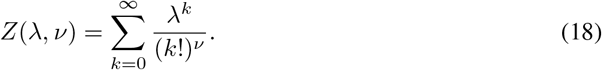

It contains the Bernoulli (*ν* → ∞), Poisson (*ν* = 1) and geometric (*ν* → 0) distributions as limiting cases. The notably property is that the CMP distribution provides a smooth transition between these well-known distributions. At integer *v*, the The moments of this distribution do not have a closed form expression in general, but can be computed using the partition function through the cumulant generating function 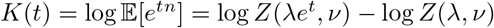. The expression for the mean and variance follow to be

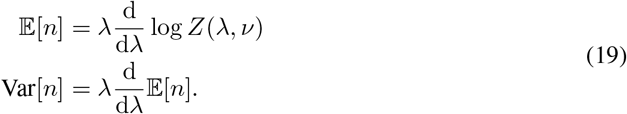

with approximate expressions [3]

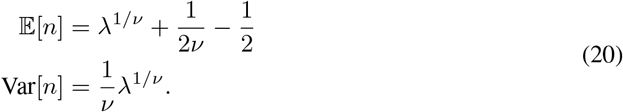

which hold well for *ν* ≈ 1 and λ > 10^*v*^.

#### A.3 Linear-softmax count distributions

The count distributions used in this work rely on a linear mapping of the input ***a*** combined with a softmax

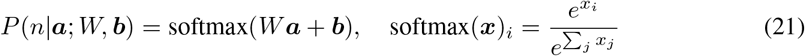

To illustrate the connection of this softmax count distribution used in Equation 1 to Poisson models, consider the distribution specified by the softmax mapping for *C* = 1 and the element-wise linear-exponential described in subsection 2.1, which in this case simply is ***ϕ***(***f***) = (*f*_1_, *e*^*f*_1_^). This choice contains the truncated Poisson distribution with *f* as the logarithm of the mean count, corresponding to *W*_*j*0_ = *j*, *W*_*j*1_ = −1 and *b_j_* = 0 with ***a*** = ***ϕ***(*f*). Hence for *C* > 1, our model is a generalization of rate-based models that implicitly assume neurons can be described by a single scalar rate parameter. The variability in such models is determined by a simple parametric relationship to the rate set by the count distribution, as can be seen for the count distribution families above.

### B. Neural dispersion and goodness-of-fit quantification

#### B.1 Kolmogorov-Smirnov framework

The measures *T*_KS_ and *T*_DS_ introduced in subsection 2.4 provide a statistical goodness-of-fit measures of the model to single neuron count statistics, and is evaluated per neuron. The predictive count distribution of the model is the reference distribution for evaluating *ξ* (Equation 7), and thus allows one to quantify dispersion *T*_DS_ (Equation 11) and goodness-of-fit *T*_KS_ (Equation 10) of the data with respect to our predictive model. By using a full predictive model, the Kolmogorov-Smirnov framework is applicable to data beyond repeatable trial structure in the inputs, such as continual recordings of freely moving animals.

For *T* values *u_t_*, *T*_KS_ is defined as

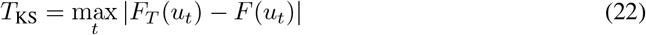

with cumulative distribution function *F*(*u*) and empirical distribution function

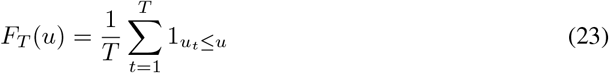

If the *u_t_* are uniformly distributed 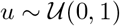 in the null hypothesis or generative model, we have *F*(*u*) = *u*. This leads to the expression given in Equation 10, which is the Kolmogorov-Smirnov statistic relevant to this work. *T*_KS_ has an asymptotic sampling distribution based on the Brownian bridge [4]. The unit Brownian bridge is defined for a Wiener process *W*(*t*) as

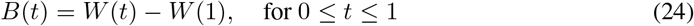

and the sampling distribution corresponds to the distribution of *N*^-1/2^ sup_*t*_ *B*(*t*).

This statistic can be interpreted as an out-of-distribution score for the observed sample, with significant misfit when *T*_KS_ is above significance value. Conventional statistics uses hypothesis testing to assess the model fit, with the null hypothesis being that the data is statistically indistinguishable from the predictive model. We can obtain model acceptance regions based on some cutoff significance value of the test statistic under its sampling distribution, often taken to be 5%. An alternative is to assess how close the empirical distribution of the test statistic is to the sampling distribution, which is the expected distribution of the statistic under the predictive model. This can be done with another Kolmogorov-Smirnov test. In this paper, we plot the acceptance regions of *T*_KS_ and show them compared to baseline models to highlight the model fit improvement on the data it was fit on. *T*_DS_ was treated similarly as a test statistic for measuring dispersion of the data with respect to the model. We present the asymptotic sampling distribution of *T*_DS_ below in subsection B.3.

#### B.2 Traditional variability measures

The traditional *Z*-score [5, 6, 7] and Fano factor [8, 2] have been used widely in the literature to quantify the variability in neural responses. The two measures are directly related

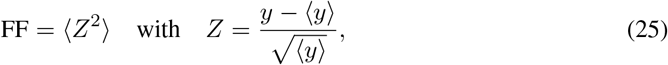

with *y* denoting spike counts and 〈·〉 the average over the relevant set of trials or time segments of experimental data. Under Poisson spiking statistics, the Fano factor is 1 and the *Z*-score is distributed as a unit normal variable. Neural data with more or less variability will lead to deviations from this reference for these dispersion measures. Activity more variable than Poisson is called overdispersed, and vice versa for underdispersed activity. These measures are mostly applied to trial-based data [9], but they can also be applied across separate time windows within a given trial or run in continual recordings. In continuous tasks as free animal navigation, the *Z*-score is often used to quantify variability or dispersion [5, 7, 1].

Note the normality of *Z*-scores under Poisson data is only asymptotically true, in the sense that we require the predicted average count 〈*y*〉 ≪ 1. The generalized *Z*-score *ξ* in Equation 7 are Gaussian under the true model by design, independent of the spike count distribution and count magnitudes. However, segments with low expected spike counts around 1 are affected significantly by the dequantization noise, hence the normality in those cases is due to the dequantization rather than model fit.

From Equation 25, we can see that our definition of a dispersion measure *T*_DS_ in Equation 11 is mathematically almost identical to the log Fano factor with *Z*-scores replaced by *ξ* (Equation 25). By using these generalized *Z*-scores, we can evaluate dispersion with respect to an arbitrary reference count distribution. However, the role of the two quantities are different. Fano factors are used to provide a measure of the spike count variability, with value 1 placing a reference point at Poisson statistics. On the other hand, *T*_DS_ is used to quantify whether the observed spike count dispersion is statistically significant compared to variability predicted by the model.

#### B.3 The sampling distribution of *T*_DS_

Under the true model, generalized *Z*-scores *ξ* (Equation 7) are i.i.d. Gaussian variables across neurons and time, hence the dispersion measure *T*_DS_ based on the sample variance of *ξ* follows a *χ*^2^-distribution. More precisely, for i.i.d. Gaussian 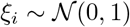, the population variance

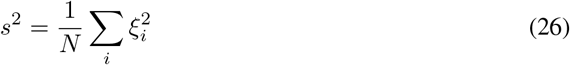

has *N s*^2^ distributed as a *χ*^2^ -distribution with *N* degrees of freedom.

The moment generating function defined as *M*(*t*) = 〈*e^-tX^*〉_*X*_ is a useful quantity for computing the moments of a distribution *p*(*X*). Note that *M*^(*n*)^ (0), indicating the n-th derivative with respect to time, gives us (−1)^*n*^〈*X^n^*〉_*X*_. When we consider the asymptotic convergence to a normal distribution of the *χ*^2^-distribution, the distribution of log s^2^ has more favourable convergence property as it is less skewed due to the logarithmic transformation [10]. Its moment generating function is

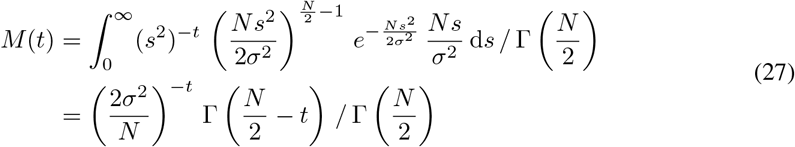

which gives rise to the cumulant function

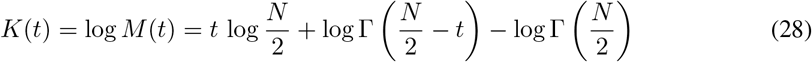

From here we can compute the first two cumulants as *κ_n_* = *K*^(*n*)^ (0) similar to the moment generating function, which are equivalent to the mean and variance of the distribution

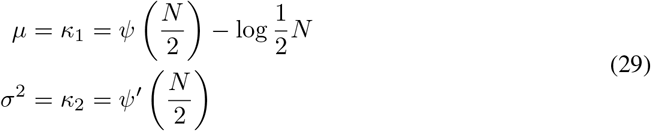

with *ψ*(*x*) = Γ′(*x*) i.e. the first derivative of the Gamma function, and the notation *f*′(*x*) = *df*(*x*)/d*x*. For values 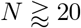, the following asymptotic expression hold well [10]

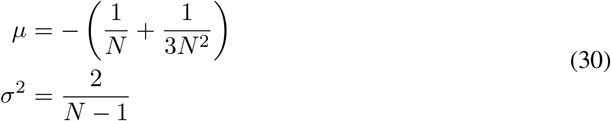

These properties lead to the construction of the *T*_DS_ metric in Equation 11. It has an asymptotically normal sampling distribution with mean 0 and variance 2/*N* – 1, convenient for statistical testing and confidence intervals.

### C Additional analysis of head direction cells

#### C.1 High-dimensional behavioural input

The universal count model (UCM) was regressed against 6 input dimensions in Figure 3A. Such high-dimensional input spaces are rife with undersampled regions, and to check if the model overfit to data we fit UCMs with a smaller number of input dimensions as shown in Figure 4A. Adding more regressors starting from only head direction progressively improves ΔcvLL, indicating the model did not overfit in the full regression case considered in this paper.

#### C.2 Temporal bin sizes

In the limit of very small time bins and *K* = 1, our model becomes a generalisation of the universal binary model [11], allowing for (observed and latent) covariates – which is critical for dissecting signal and noise correlations in neural data. In fact at *K* = 1, the distinction between classical models assuming Poisson variability and our universal model allowing for non-Poisson variability becomes irrelevant as all possible spike count distributions (SCDs) in a time bin are Bernoulli. Thus, in this limit, our model also becomes conceptually very similar to GLMs [12] and GPFA [13] (with binary emissions) in that conditioned on covariates (potentially including the spiking history of a neuron itself, or of other neurons, in GLMs) spiking becomes an inhomogeneous Poisson process. However, these previous models remain Poisson-like *at all time scales* (in the sense that, conditioned on covariates, spike counts remain Poisson distributed) and only allow covariates to modulate the mean firing rate. This is because, given covariates, spiking is assumed to be independent in consecutive infinitesimal time bins, a key property of the Poisson point process. The key advance of our work is precisely that *even at larger time bins* (and with *K* > 1) it is not restricted to Poisson count distributions and allows covariates to modulate any spike count statistic. Indeed, we found that experimental data deviated from Poisson-like statistics in important ways and was in many cases substantially modulated by covariates at the 40 ms time bin size (with maximum spike count K =11) that we chose (Figure 3B and C).

The advantage of choosing essentially infinitesimally small time bins is that there is no “arbitrary” (though see above) time bin size parameter and every individual spike can be predicted (at least in principle, see also note below). The disadvantage is that, as we explain above, in this case it is not even conceptually possible to model modulations of response variability independent from modulations of firing rates, as the two are inseparable in the underlying Bernoulli model (as K =1). Indeed, experimental studies of neural variability, and its modulation by covariates, have always used larger time bins to compute Fano factors or *Z*-scores [14, 1, 5, 9]. Our work is a direct generalization of this perspective, extending beyond rigid trial-structure. At the other extreme, the disadvantage of choosing time bins that are too large is that it may miss the time scale at which covariates modulate neural firing. In particular, when studying phenomena at time scales comparable to the interspike intervals, such as theta phase precession [15, 16], binning may average away such effects if the bin size is too large. However, binning does reduce the number of total time points and is thus more practical for studying the activity of large populations recorded over long time periods.

Our choice of 40 ms time bins was based on previous empirical measures of autocorrelation time scales in neural activity [17, 18]. We also ensured none of our behavioural covariates had a shorter time scale (see Figure 3G). In Figure 4B, we can see the sensitivity of the count analysis to bin size. We fit separate UCMs to data at various bin sizes as presented in the figure. Note that increasing the bin size leads to higher Fano factors. In addition, notice the consistent decrease of variability at higher firing rates. Temporal correlations in the spike trains generally lead to spike counts being correlated across consecutive time bins. This in turn leads to potentially more extreme fluctuations in the sums of consecutive counts (qualitatively identical to picking a larger bin size), and thus higher or lower variability. The exact details depend on the underlying process, and in general no analytical treatment is possible. For stationary renewal processes however, an analytical treatment of Fano factor dependence on bin size is available [19].

In summary, there is nothing to say *a priori* that *the right* time bin size for studying neural variability is the infinitesimally small limit used by several previous approaches. Indeed, our empirical results showing non-Poisson conditioned SCDs at longer time scales suggest that longer time scales may be more appropriate – at least in the data set we analysed. In general, we argue (see above) that if one wants to study the modulation of neural response variability then one must use appropriately sized (non-infinitesimally small) time bins. In turn, in this setting, our approach is unique in offering a statistically principled method to do so and offers novel insights into the variability of head direction cells in mice.

#### C.3 Drifting and ATIs

Joint tuning curves can reveal neural representations that are not factorized over a set of covariates. The Bayesian nature of Gaussian processes takes care of undersampled regions that are rife in high dimensional input spaces, which is the setting in this work for studying joint tuning to high-dimensional behavioural input. By looking at joint tuning curves between particular covariates, the model reveals properties that have been observed in separate experimental works.

One can pick up representational drift [20, 21] by regressing against absolute time of the recording. We can then compute a linear drift by quantifying how much the preferred head direction θ_pref_ (Equation 40) changes with time using circular linear regression (subsection D.8). Not all cells are well-described by a linear drift in time, and only cells that have a sufficiently good fit to the regression line are included. The time regressor needs to be considered carefully as it may confound at time scales of latent trajectories. As long as the time scale in the kernel is much larger than the time scale over which the latent variables vary, we can interpret the temporal drift as a separate process from the latent trajectories. Indeed, by initializing at time scales equal to half the total recording time, these time scales of the Gaussian process kernel remain significantly higher than any behavioural time scale (Figure 3G). We find most cells cluster at a drift of ≈ 20 °/hr.

The joint AHV-HD plot reveals anticipatory tuning: when animals turn their head, the head direction tuning curves shift in response to head rotations such that cells expected to spike appear to fire earlier than expected. Theoretical studies have shown that this improves temporal decoding, in the sense that the bias-variance trade-off for decoding downstream can be improved with anticipatory tuning [22]. It appears that the head direction population anticipates the future head direction based on current movement statistics, which allows one to reduce the bias introduced with causal decoding. One can define an anticipatory time interval (ATI) analogous to linear drift, as the amount of change in preferred head direction with angular head velocity. This is again quantified using circular regression (subsection D.8). Similarly, not all cells are well-described by a linear relation between *θ*_pref_ shift and angular head velocity *ω*, and only cells that have a sufficiently good fit to the regression line are included. Note the ATI values in Figure 4C are negative, while in the literature they are postive and differ per region. The neural data description files [23] did mention that the zero time frame of behaviour was randomly misaligned to neural spiking data up to 60 ms. Behaviour may be shifted with respect to the neural spike train, indeed in preliminary analyses with shifted spike trains we found values consistent with literature for ATIs when shifiting ≈ 60 ms [22].

#### C.4 Latent variable analysis of head direction data

In Figure 4D, we see the inferred angular latent signal is closely related to the head direction. This is similar to previous analyses with non-Euclidean Gaussian process latent variable models [24] and presents an exceptional case where the inferred latent signal is directly relatable to an experimentally observed variable. Therefore, a measure of error can be computed between the two. To do so, we align the latent signal to the observed head direction by fitting a transformation of the form described in Equation 45. Latent trajectory root-mean-squared error (RMSE) was computed with 3-fold cross-validation, using the geodesic on the ring. We align the latent trajectory Equation 45 to the behaviour in the fitting segment, and compute the geodesic RMSE on the held-out validation segment (see subsection D.8).

The UCM again shows improvement over baseline models, and interestingly the inferred latent signal is more correlated to the behavioural head direction (left bottom panel of Figure 4D). When shifting the behaviour w.r.t. the latent signal in the UCM model, we observe a minimum cross-validated RMSE at around −100 ms. We thus tentatively identify a delay in the signal represented compared to measured behaviour, with the behaviour lagging the latent signal. From the UCM fit at zero behavioural shift, the inferred linear drift compared to observed head direction is 3.1 ± 1.2 °/hr, see right top panel of Figure 4D. This is smaller but in the same direction as the drift found using regression in the tuning curves in panel C, which cluster around 20 °/hr.

### D Analysis details

#### D.1 Synthetic data

We construct a synthetic head direction cell population inspired by bump attractor models [25, 26, 24]. Firing rate tuning curves to head direction *θ* are parameterized as von Mises bump functions with some constant offset

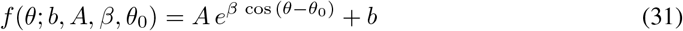

with *b* > 0 and *A* > 0. This results in *f* ≥ 0 for all valid inputs and parameters. For modelling firing rates, we additionally restrict ourselves to *β* > 0 to avoid inverted bumps at the preferred head direction *θ*_0_.

In the Conway-Maxwell-Poisson (CMP) synthetic population, we placed the tuning curves from Equation 31 on parameters *ν* and the approximate mean 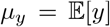 in Equation 20. Note both parameters have to be non-negative to be valid. Furthermore, the tuning curves of *ν* had potentially negative *β* ∈ ℝ and different parameter statistics than for *μ_y_*. Again, these were chosen such that firing rates and variability were within the physiological regime. To roughly match the mean counts with von Mises bump pattern amplitudes, we used the mapping

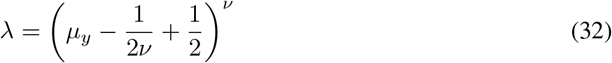

which was based on the approximate relation Equation 20 of the mean. We chose a time bin of 100 ms, which led to *K* = 18 in the synthetic data generated. To sample from the CMP distribution once we specified λ and *ν*, we used the fast rejection sampling method [27].

For the modulation by a hidden Euclidean signal in the modulated Poisson population, we additionally placed Gaussian tuning curves on the latent dimensions with varying standard deviations and means. The Gaussian tuning curves tiled the latent space that was traversed, which allowed the model to infer the full trajectory. Note that tuning is factorized across the two dimensions (head direction x and latent signal z). Parameters were randomly sampled from distributions that led to firing rates and variability within the physiological regime. We again picked a 100 ms time bin, which gave *K* = 28.

#### D.2 Neural data

Data was taken from Mouse 28, session 140313, during the wake phase [23]. The spiking data was recorded at a resolution of 20000 Hz, whereas behaviour was extracted from video recordings of animal body tracking at a resolution of 39.06 Hz. Note the time of the first video frame was randomly misaligned by 0–60 ms to the neural spike trains. We removed invalid behavioural segments in the data and performed linear interpolation across those segments. For circular variables, interpolation was taken in the shortest geodesic distance. We binned spiking data at 1 ms, and interpolated behavioural data to reach the same sampling frequency that is higher than the behavioural recording frequency. At a binning of 40 ms used in our analysis, we had *K* = 11 as the maximum count value.

We selected head direction cells based on a sparsity criterion, after trying several criteria as mutual information typically used for place cells [28]. First, we binned the head direction variable into 60 equal bins over the range [0,2*π*]. For each bin, we now compute the average spike counts *y_i_* for head directions within bin *i*, and the relative occupancy *P_i_*. Note ∑_*i*_ *P_i_* = 1 is a probability distribution. Sparsity is defined as

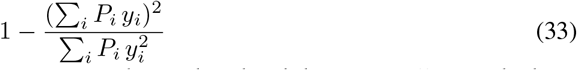

and with a selection criterion of sparsity ≥ 0.2 we obtained 33 head direction cells, of which 15 are in postsubiculum. Alternatively, although more computationally intensive, we could directly regress a Gaussian process model (e.g. Poisson baseline model Equation 34) and look at the kernel lengthscales on the angular input dimension. These will be appreciably larger than 2*π* for cells that are not tuned much to head direction.

Note that quite a few head direction units, which are supposed to represent single cells, show bimodal tuning curves or more to head direction. This is likely due to multiple neurons as signals can pollute in electrophysiological recordings and spike sorting can fail to distinguish between them [29, 30].

#### D.3 Baseline models

##### D.3.1 Gaussian process models

The log Cox Gaussian process model puts a GP prior on the rate function of an inhomogeneous Poisson process (Equation 13) with an inverse link function *f*(*x*) = *e^x^* that is exponential

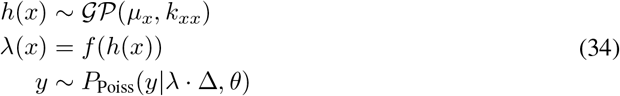

where the time bin length is Δ, which turns λ into a proper rate quantity.

The heteroscedastic negative binomial model builds on this encoding model, More precisely, two GPs with an exponential inverse link function are used to model tuning to covariates of the rate λ and inverse shape 1/*r* of the negative binomial likelihood (Equation 16), leading to the model

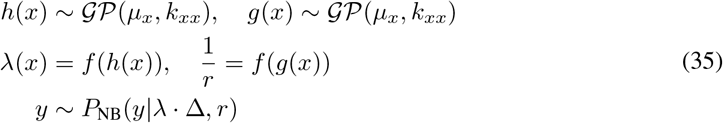

In the same spirit, we could construct the more flexible heteroscedastic Conway-Maxwell-Poisson model. This model would be able to capture both over- and underdispersed count data (Fano factors above and below 1), but it has difficulty in scaling to large data due to the series approximation of the partition function in Equation 17.

##### D.3.2 Artificial neural network models

The artificial neural network (ANN) model used to replace the Gaussian process in validation experiments was designed such that there was sufficient expressivity to model the neural activity. In fact, we see that the neural network overfits in Figure 2A, which indicates that there was enough capacity in the network. The network architecture consists of an input layer providing ***x***_*t*_ (and latent ***z***_*t*_ when present), encoding angular dimensions *θ* as a two-dimensional vector (cos *θ*, sin *θ*). There are 3 hidden layers containing 50, 50 and 100 hidden units in order from input to output layer, with sinusoidal activation functions to construct smooth overall mappings [31]. The output layer consisted of *N* · *C* linear units providing *f_cnt_* in Equation 1, with *N* the number of neurons and *C* the number of degrees of freedom per neuron (which was 3 in this work). The UCM with an ANN mapping leads to a model similar to VAEs [32] with a softmax likelihood and free variational parameters instead of amortization with an inference network.

#### D.4 Computing generalized *Z*-scores

The generalized *Z*-scores *ξ* in Equation 7 provide a normalized quantification of neural activity under the predictive model. For UCM, the count distribution *P*(*y*) which is used to compute is taken to be the mean posterior count distribution of the posterior *q*(Π|*X, Z*). In the case of baseline models, the reference *P*(*y*) is given by the parametric distribution (Poisson in Equation 13, negative binomial in Equation 16) evaluated at the mean posterior values of the count distribution parameters given by the Gaussian process mapping (see Equation 34 and Equation 35). This is strictly speaking different from the mean posterior count distribution, as the parametric distribution depends nonlinearly on these parameters. However, the difference is insignificant when the variational uncertainties are small, which was the case in practice.

#### D.5 Marginal and conditional tuning curves

Due to the high dimensional input space, we can either visualize slices of the tuning curve over the relevant input variables ***x***\* or instead marginalize over other input variables. The conditional tuning curves are based on the count distributions *P*(*y*|***x***\*, ***x***^*c*^), where ***x***^*c*^ are fixed and cover the dimensions complementary to ***x***\* (these are plotted in Figure 3C). On the other hand, marginalizing over ***x***^*c*^ depends on the chosen *p*(***x***^*c*^). A natural perspective is to consider the input data distribution 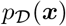 and treat the observed input time series as a Markov Chain Monte Carlo path sampled from it. We can use this to approximate the exact marginalization, which is intractable as we do not know 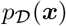. Conceptually, this is equivalent to an experimenter only looking at neural tuning to ***x***\*, which automatically marginalizes over all other behaviour not included during the experiment. We denote observed input with a subscript 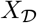 in this scenario, to distinguish it from chosen input locations. We only consider the ***x***^*c*^-dimensions of the joint density 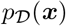 and this mathematically becomes

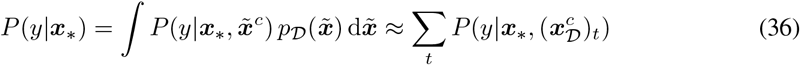

which defines the marginalization through the computation done in practice (summing over the time series of observed ***x***^*c*^ while keeping ***x***\* fixed). From this marginalized distribution, we can compute similarly quantities like the mean spike count or count variance.

For the tuning indices, we evaluate the the count statistic *T_y_*(***x***\*) with respect to the posterior mean distribution *P*(*y*|***x***\*) after marginalizing (order does not matter as both are sums) to compute the tuning indices as described in Equation 6. Optimization over ***x***\* of *T_y_* (***x***\*) is done by grid search, as ***x***\* is low-dimensional and we compute its values over a grid anyway for plotting tuning curves of mean, Fano factor or any other count statistic.

We used 300 Monte Carlo samples from *q*(Π|*X, Z*) to compute the conditional tuning curves plotted in this paper. For marginalized tuning curves, we use 100 MC samples and temporally subsampled the observed input XD to retain the first time step per every 10 time steps, and used this to evaluate Equation 36. As behaviour shows strong temporal correlations at short time scales (Figure 3G), this allows us to estimate the marginal tuning curves more efficiently. The mean of these samples was used to compute the mean posterior tuning curves for evaluating the TIs. When evaluating the average mean count and Fano factor at every time step (Figure 3B and Figure 4B), we used 10 MC samples from *q*(Π|*X, Z*). When latent variables were present (Figure 3E), the 10 MC samples were drawn from *q*(*Z*), corresponding to *m* = 10 and *k* =1 in Algorithm 1.

#### D.6 Temporal cross-correlations of covariates

We use the cross-correlation between time series *x_t_* and *y_t_*

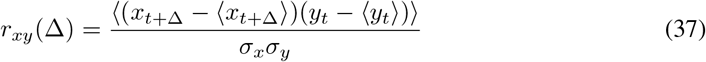

which includes the auto-correlation as a special case, e.g. *r_xx_*(Δ). When one of the variables is a circular variable *θ_t_*, we use the linear-circular correlation coefficient in [33]

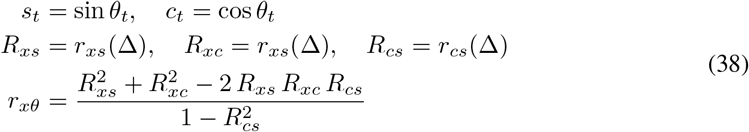

and for the case when both are circular, we use the circular correlation coefficient proposed by [34]

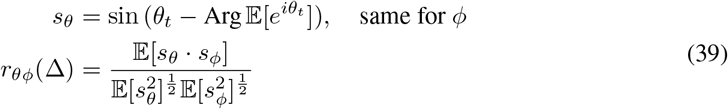

Time scales are estimated from the auto-correlations of covariates. The time scale *τ* is then chosen as the time step at which the value of the auto-correlation dropped by a factor *e* from 1 at Δ = 0.

#### D.7 Preferred head direction

To compute the preferred head direction *θ*_pref_, we use the centre-of-mass of the firing rate profile *r*(*θ*) of head direction *θ*

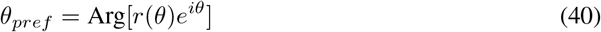

which is more robust to noise than taking the angle at which *r*(*θ*) is at a maximum. We can evaluate *θ*_pref_ as a function of angular head velocity (AHV) and absolute time to compute the ATIs and the neural drift as described in subsection C.3.

#### D.8 Circular-linear regression

We computed the circular-linear regression [35] using a measure of the correlation between circular variables *θ*_1_ and *θ*_2_

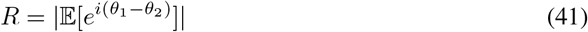

By computing *R* between a circular-linear function *ε*(*t*)

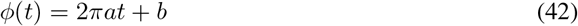

and the circular data time series *θ_t_*, we can perform the regression by maximizing *R* through optimizing the parameter *a* with gradient descent. The offset *b* is obtained analytically

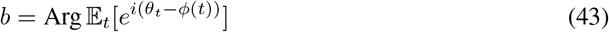

From the values *a* after fitting, one can compute the linear drift values and ATIs as described in subsection C.3. In addition, not all cells are well-described by the linear drift or ATIs, so we discarded cells which had an optimized value of *R* < 0.999. This cutoff was chosen as it retains cells that are visually in agreement with linear relations as seen in Figure 4, while discarding a few outlier cells.

#### D.9 Latent alignments

To align 1D circular latent trajectories *z_c_* to a target trajectory, we minimize their mean geodesic distance under a constant shift *μ* and potential sign flip *s* = ±1

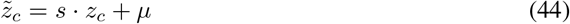

We add a linear drift Δ to find potential drifting of the inferred trajectory

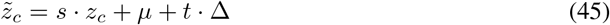

as done in panel D of Figure 4. This is similar to the circular-linear regression above [35], but with the geodesic distance on the ring instead. This is consistent with root-mean-square errors in the latent signal from behaviour that are computed with the geodesic distances. For 1D Euclidean latent trajectories, we align by fitting a translation and scaling parameter.

In all cases, the root mean squared error (RMSE) of the alignment is evaluated in a cross-validated manner. For circular variables, we use the geodesic distance for computing the squared error just as in aligning. In more detail, we fit the trajectory transformation parameters such that we minimize the errors on the validation segment, and then use these fitted parameters to compute the transformed latent trajectory in the held-out segment. This is then used to compute the RMSE for the alignment of the cross-validation fold.

### E Implementation details

#### E.1 Mathematical details of the optimization objective

##### E.1.1 The sparse Gaussian process posterior

Exact Gaussian processes (GPs) have *O*(*T*^3^) computational complexity and *O*(*T*^2^) memory storage with *T* input points [36]. This is unfavourable for scaling to large or massive datasets. In addition, non-Gaussian likelihoods lead to intractable marginal likelihoods and hence one needs an approximate optimization objective. Stochastic variational inference [37] provides a framework for applying Gaussian process methods using non-Gaussian likelihoods and approximations for scalability. Let us denote the exact GP prior by *p*(***f***) with vector ***f*** the latent function points at the input locations *X*, the likelihood by *p*(***y***|***f***) with observed data ***y***, and the approximate posterior by *q*(***f***). One then needs to be able to (1) efficiently and differentiably sample from *q*(***f***), and (2) efficiently evaluate and differentiate the Kullback-Leibler (KL) divergence between *q*(***f***) and *p*(***f***).

Sparse approximations [38] reduce the computational complexity to *O*(*MT*^2^ + *M*^3^) and storage to *O*(*M*^2^) with *M* inducing points, which effectively aim to summarize the input data with a smaller set of points. Such methods are scalable for large *T* as long as *M* ≪ *T* provides sufficient modelling flexibility. The key idea is to extend the function values ***f*** with additional function values ***f***_*u*_ at inducing points. Let us denote inducing points with function values ***f***_*u*_ at inducing point locations *U*, which we jointly learn with other variational parameters. The GP kernel evaluated at function point locations is denoted by *K_XX_*, and at inducing point locations by *K_UU_*. Cross-covariances are denoted by *K_XU_* and *K_UX_*. The joint variational distribution to the augmented Gaussian process posterior *p*(***f***, ***f***_*u*_|***y***) is defined as

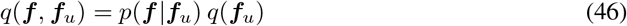

where the variational distribution 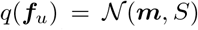, and *p*(***f*** |***f***_*u*_) is the conditional Gaussian distribution from the generative model. The variational distribution over GP function values *q*(***f***) is simply obtained by marginalizing out ***f***_*u*_, which leads to a Gaussian with

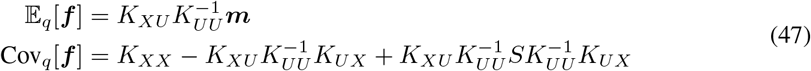

and the KL divergence

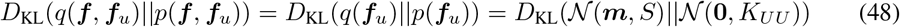

which can be evaluated as long as *M* is not too large. The reason for the choice in Equation 46 becomes clear: due to the cancellation of *p*(***f*** |***f***_*u*_), we do not have to invert large matrices related to *K_XX_*. Note that sampling from *q*(***f***) for a large input set X is problematic [39]. Fortunately, our likelihood factorizes across time and thus we can evaluate the expectation under *q*(***f***) with the diagonalized distribution for which sampling is trivial.

Unlike purely variational approaches, the approximate posterior in Equation 46 amortizes the inference through the learned inducing points, and allows one to obtain a predictive distribution using the approximate posterior evaluated at a new set of inputs *X*\*

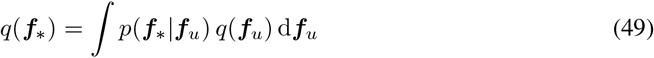

This property also allows one to apply mini-batching or subsampling to Gaussian processes [40, 41]. Overall, this leads to the Sparse Variational Gaussian Process (SVGP), combinining Sparse Gaussian Processes [38] with stochastic variational inference [32].

To accelerate convergence, a different parameterization of the variational distribution was used. One performs a change of variables 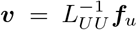 with 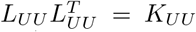 from the Cholesky decomposition. This transforms *p*(***f***_*u*_) into 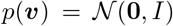, referred to as whitening, and the variational parameters are now defined for 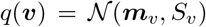 [41]. In practice, matrix-vector products with 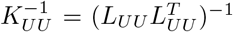 are evaluated by solving two triangular systems with *L_UU_*. The whitened representation simplifies Equation 47 as we do not need to compute 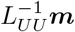 and 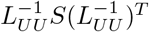 in the non-whitened parameterization. The KL divergence Equation 48 also simplifies as we now have unit normal *p*(***v***).

To increase the expressivity of multi-output GPs, a separate set of inducing points locations is used for each output dimension (neuron in this work), along with separate kernel hyperparameters as lengthscales for each input and output dimension. This is equivalent to modelling each output dimension by a separate GP, and leads to an overall computational complexity of *O*(*NCTM*^2^) and storage of *O*(*NCM*^2^) for our model (see section 2 for definition of quantities). A thorough description of a scalable multi-output SVGP framework is given in [42]. We define the multi-output variational posterior *q*(*F*) as

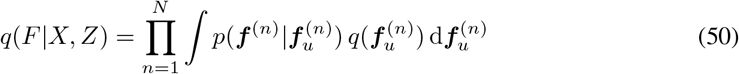

with output function values *F* evaluated at input locations *X, Z*.

##### E.1.2 Generative model and variational inference

The overall generative model Equation 1 as depicted in Figure 1 is

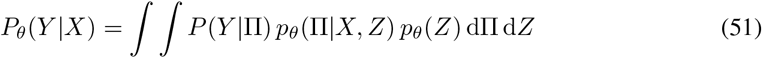

with the product of individual count distributions *P*(*Y*|Π). The model parameters *θ* include the GP *θ*^GP^ and the prior *θ*^pr^ (hyper)parameters, as well as the softmax mapping weights *W_n_* and biases ***b***_*n*_. Note that the distribution over count probabilities

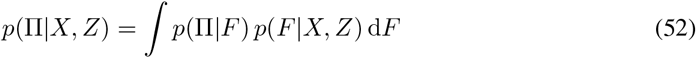

contains the Gaussian process prior *p*(*F*|*X, Z*) over *F*. The mapping from *F* to Π denoted by Π(*F*) (Equation 1) is deterministic, and therefore *p*(Π|*F*) is a delta distribution *δ*(Π - Π(*F*)).

The exact Bayesian posterior over Π and *Z* is intractable, hence we use an approximate posterior as defined in Equation 3. The variational parameters *φ* specify the latent variational posterior, while *χ_u_* consists of inducing point locations *X_u_* and the means and covariance matrices of *q*(*U*) for the sparse Gaussian process posterior *q*(*F*|*X, Z*) (Equation 46). The wrapped normal distribution used for circular dimensions in *q*(*Z*), i.e. dimensions with *z* ∈ [0, 2*π*), takes the form [43]

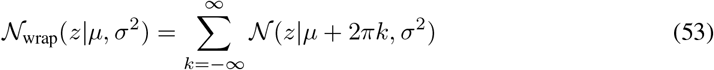

and was evaluated with a finite cutoff at *k* = ±5 of the infinite sum. This is an accurate approximation as long as *σ* ≪ 2*π*. When plotting the standard deviations of the approximate posterior *q*(*Z*), we plot *σ* for both Euclidean as well as circular variables. This is similarly an accurate approximation in the circular case when *σ* ≪ 2*π*, which was true in practice.

The marginal likelihood in Equation 51 is intractable. Instead, we minimize the negative ELBO or variational free energy loss objective using our approximate posterior

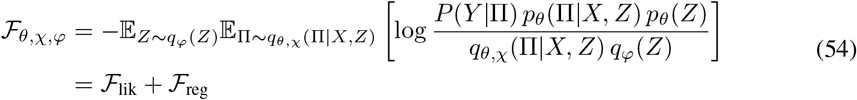

which is an upper bound to the negative log marginal likelihood [32, 41]. The objective decomposes into a log likelihood expectation term 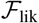 and some regularization terms arising from the model priors 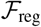. These terms are amenable to Monte Carlo evaluation or quadrature approximation as we show next, and in some cases are even available in closed form.

The variational expectation of the log likelihood

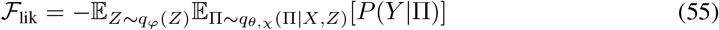

can be evaluated using Monte Carlo sampling to obtain unbiased estimates in the general case. As an alternative method, Gauss-Hermite quadratures can provide a deterministic approximation to the expectation with respect to *q*(*F*|*X, Z*) [41]

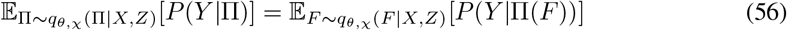

where Π(*F*) denotes the transformation from *F* to count probabilities as in Equation 1. Here we used

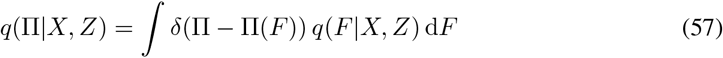

analogous to Equation 52 for the generative model. This corresponds to a zero variance estimator with a small bias for sufficiently many quadrature points. As the likelihood factorizes over time, the expectation with respect to the multivariate variational posterior *q*(*F*) factorizes into expectation terms with univariate Gaussian distributions, their variances taken from the diagonal of the covariance matrix. Because of this, we only need MC sampling from univariate distributions, allowing us to work with many time points and large batch sizes. This removes correlations between posterior function sample points at different input values, which cannot be done when factorization over time does not hold (e.g. GP priors in deep GPs). The issue of efficiently sampling from the full posterior has been considered in [39].

The regularization terms can be written as Kullback-Leibler divergences

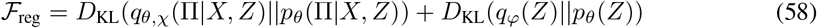

with the ratio of *q_θ_*(Π|*X, Z*) and *p_θ,χ_*(Π|*X, Z*) in the first KL divergence equivalent to

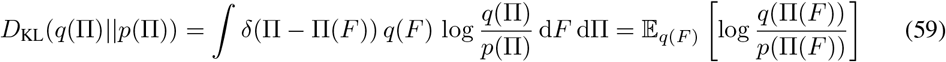

where we have made the mapping Π(*F*) explicit. In practice, the choice of *C* < *K* implies this mapping is underparameterized and generally injective for matrices *W* of rank ≥ *C*. When we are in the universal limit *C* = *K* and *W* is rank *K*, the mapping will be bijective. As long as the mapping from *F* to Π is not many-to-one, we have the continuous random variable transform

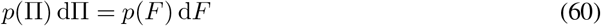

and this leads to Equation 59 becoming *D*_KL_(*q*(*F*)||*p*(*F*)). Hence, the regularization terms in the loss objective Equation 54 consist of KL divergences

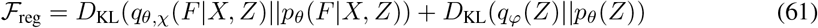

that can be computed with analytical expressions in the case when all distributions are Gaussian.

#### E.2 Latent space priors

We use the Markovian priors as specified in Equation 2, and these priors can be specified on different manifolds [24]. For Euclidean spaces, we use the linear dynamical system prior

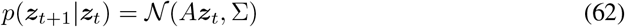

In particular, we use diagonal Σ and *A* to learn factorized latent states. We constrain *A_ii_* = *a_i_* ∈ (− 1,1) for stability, and we fix 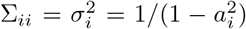 to obtain a prior process with stationary variance 1 while optimizing for *a_ii_*. On the toroidal manifold, we use

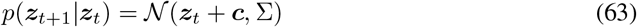

as due to rotational symmetry *A* = *I*. Again, we use diagonal Σ. Both ***c*** and 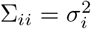 are learned as part of the generative model.

When temporally batching input, one has to be careful to retain the continuity in the prior *p*(*Z*) with the previous batch (beyond the first batch at the start). This is done by ensuring that the first ***z***_*t*_ in the batch is the last step in the previous batch, and this will correctly subsample the prior *p*(*Z*) defined over the entire input time series. When performing cross-validation with validation segments within the overall input time series, we treat the gap as a discontinuity in the latent trajectory and do not include the latent state right before the validation segment.

#### E.3 Gaussian process kernel functions

In this work, we used the RBF kernel defined on Euclidean and toroidal manifolds [24]. In particular, this kernel function is given by

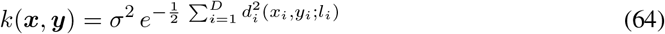

with rescaled distances

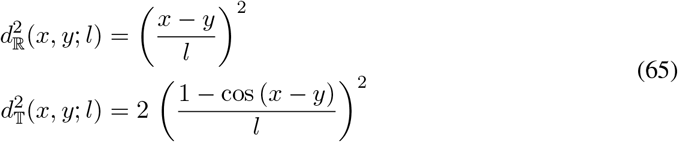

for Euclidean and toroidal spaces ℝ and 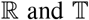, respectively. To cover different input dimensions of different topologies, we use product kernels with suitable distances *d* per input dimension, resulting in sums over dimensions in Equation 64. These distance functions can be used to extend other kernels such as Matérn kernels to non-Euclidean spaces [24].

#### E.4 Overall algorithm and code

The outline of the inference procedure is given in Algorithm 1. Additionally, instead of drawing Monte Carlo samples for the Gaussian variational posterior *q*(*F*), we provide the option to compute the Gaussian expectation using Gauss-Hermite quadratures [41]. This was used to estimate the cvLLs (Equation 9) for models after training, which reduced stochasticity in the cvLL estimate with a negligible bias when using 100 quadrature points. Monte Carlo samples or quadrature point dimensions are parallelized over in addition to other dimensions like neurons or time, using extra tensor dimensions in modern automatic differentiation libraries. We use PyTorch [44] to implement the algorithm for inference of our models. For optimization, we use Adam [45] with no weight decay and default optimizer hyperparameters in PyTorch.

The code provided^2^ contains a library under the name ‘neuroprob’, which was written to organize the implementations of Gaussian process and GLM based models with different likelihoods as used for baseline models in this paper. In addition to count likelihoods, it contains an implementation of spike-spike and spike-history couplings [12, 46] and modulated renewal processes [47, 48] to deal with data at the individual spike time level. All models can be run with both observed and latent inputs on Euclidean and toroidal manifolds [24].

**Algorithm 1.**
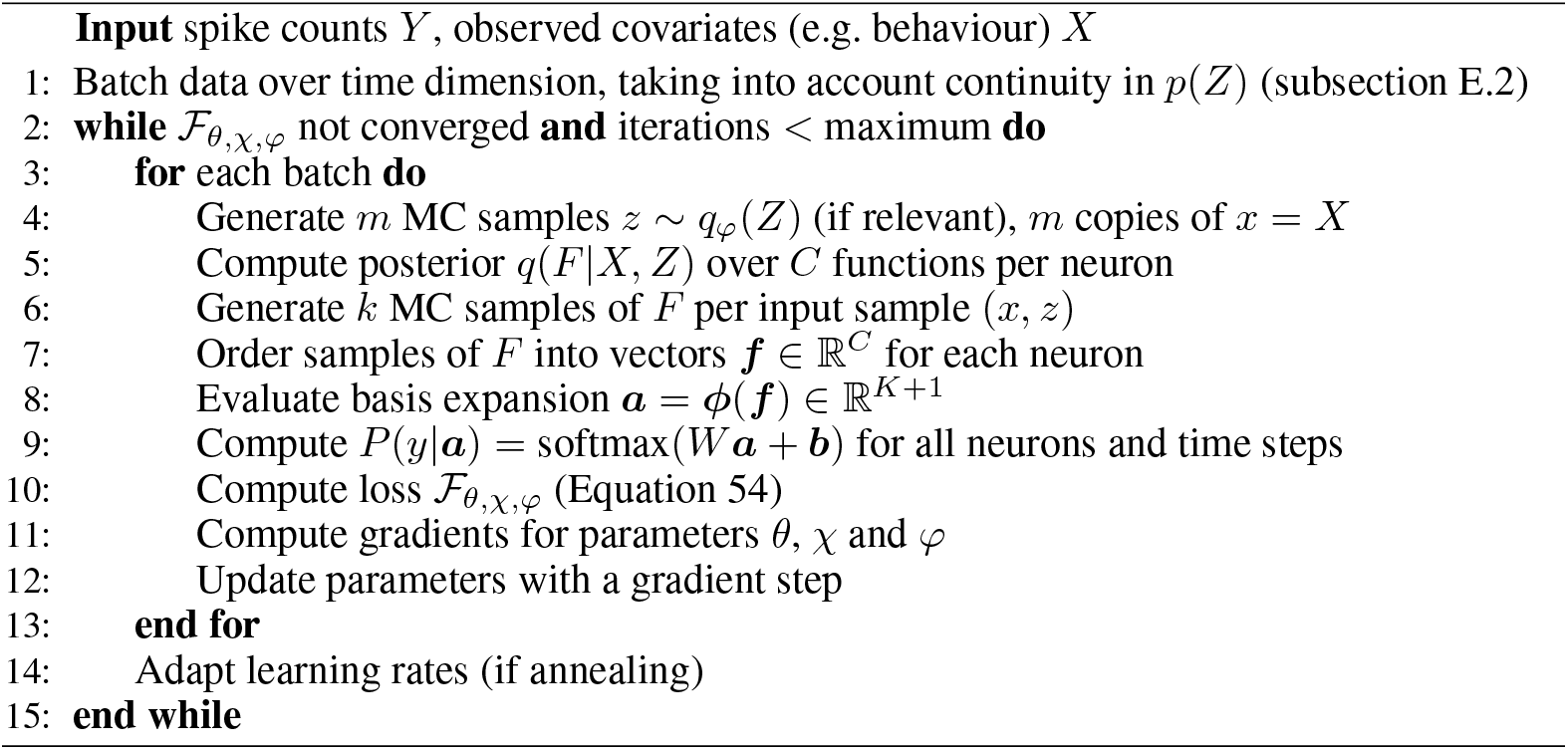
Joint latent-observed input inference scheme

#### E.5 Model fitting

##### E.5.1 Inducing point initialization

The first input dimension had its inducing points uniformly spaced between 0 and 2*π* for circular dimensions, and −1 to 1 for Euclidean latent dimensions. Observed dimensions had natural intervals defined by the behavioural statistics (e.g. 0 to the mean animal speed), and we placed inducing points uniformly throughout this interval. For the other dimensions, we initialized random inducing point locations based on the topology of the input variable. We place Euclidean variables as a random uniform distribution in its corresponding interval as described previously, while circular variables took on random uniform values in [0,2*π*].

The number of inducing points has been shown to scale favourably as *O*((log *T*)^*D*^) for standard Gaussian process regression models [49]. In this work, we used *O*(*D* log *T*) which captured rich tuning and satisfactory model fits combined with the flexible count distributions. The suggested *O*((log *T*)^*D*^) does become computationally expensive for high dimensional input, and was not tried with the high-dimensional regression models.

##### E.5.2 Fitting details

We select the model with the lowest loss from 3 separate model fits, initialized with randomized inducing points as described above. The maximum number of training epochs was 3000, but we stopped training before if the loss did not decrease more than ≈ 10^-3^ percent over 100 steps. The learning rate was set to 10^-2^, and we also anneal the learning rate every 100 steps by a factor 0.9. This was for both Gaussian process as well as artificial neural network models. In the case of latent spaces, we used a learning rate of 10^-3^ for standard deviations of the variational distribution *q*(*Z*). All cases lead to satisfactory convergence of the model.

For latent variable models with a single angular latent, we initialize the lengthscale at large values. This avoided the model to overfit and fold the latent space as seen in panel A of Figure 3 for the ANN model. For these models, the best fits were achieved with an initial learning rate of 3 · 10^-2^ and 5 · 10^-3^ for the kernel lengthscale and the standard deviations of the variational distribution *q*(*Z*).

##### E.5.3 Hardware and fitting time

Synthetic data was analyzed with GeForce RTX 2070 (8 GB of memory). Real data was analyzed with Nvidia GeForce RTX 2080Ti GPUs (with 11 GB of memory). Fitting 33 neurons with ~ 6 · 10^4^ time points with the regression model in Figure 3 takes around 20 minutes, while fitting with a four dimensional latent spaces added takes around 50 minutes. These numbers can fluctuate depending on the flexible stopping criterion above. Generally, there is a trade-off between memory usage and speed by setting the batch size, with larger batch sizes being generally faster but taking more memory.

## Footnotes

1 For notational convenience, *X* denotes both observed and latent covariates here.

2 https://github.com/davindicode/universal_count_model

